# Tertiary lymphoid tissue develops during normal aging in mice and humans

**DOI:** 10.1101/749200

**Authors:** Marianne M. Ligon, Caihong Wang, Zoe Jennings, Christian Schulz, Erica N. DeJong, Jerry L. Lowder, Dawn M. E. Bowdish, Indira U. Mysorekar

## Abstract

Aging has multifaceted effects on the immune system in the context of systemic responses to specific vaccines and pathogens, but how aging affects tissue-specific immunity is not well-defined. Chronic bladder inflammation is highly prevalent in older women, but mechanisms by which aging promotes these pathologies remain unknown. Here we report distinct, age-associated changes to the immune compartment in the otherwise normal female (but not in male) mouse urinary bladder and parallel changes in older women with chronic bladder inflammation. In aged mice, the bladder epithelium became more permeable, and the homeostatic immune landscape shifted from a limited, innate immune-predominant surveillance to an inflammatory, adaptive immune-predominant environment. Strikingly, lymphoid cells were organized into tertiary lymphoid tissues, hereafter named bladder tertiary lymphoid tissue (bTLT). Analogous bTLTs were found in older women, many of whom had a history of recurrent urinary tract infection (UTI). Aged mice responded poorly to experimental UTI, experiencing spontaneous recurrences at higher rates than young mice. However, bTLT formation was dependent on aging and independent of infection. Furthermore, bTLTs in aged mice played a role in *de novo* antibody responses and urinary IgA production by recruitment of naive B cells that form germinal centers and mature into IgA-secreting plasma cells. Finally, TNFα was a key driver of bTLT formation, as aged TNFα^-/-^ mice lacked bTLTs. Both aged TNFα^-/-^ and wild type mice exhibited increased bladder permeability, suggesting that epithelial dysfunction may be an upstream mediator of chronic, age-associated bladder inflammation. Thus, bTLTs arise as a function of age and may underlie chronic, age-associated bladder inflammation. Our model establishes a platform for further investigation of age-association tissue inflammation and translation to new treatment strategies.

**One Sentence Summary:** Mice develop bladder tertiary lymphoid tissue (bTLT) during aging that is dependent on TNFα and independent of urinary tract infection.

## INTRODUCTION

Immune dysfunction during aging is characterized by chronic, low-grade inflammation coupled with ineffective responses to pathogens. Aging is also the strongest risk-factor for numerous chronic diseases, including cardiovascular disease, neurodegeneration, osteoarthritis, and cancers. While these diseases are all linked by chronic inflammation, immune responses vary by tissue, resulting in tissue-specific inflammation and dysfunction. Older women (50+) are highly susceptible to bladder disorders including overactive bladder/urge incontinence (OAB), interstitial cystitis/bladder pain syndrome (IC/BPS), and recurrent urinary tract infections (rUTIs) (*1-5*). These disorders all have a chronic inflammatory component as well as overlapping symptoms, known as lower urinary tract symptoms (*6, 7*). How and why bladder disorders and bladder inflammation become more prevalent with aging is not currently understood (*8*). Since aging is characterized by chronic, low-grade systemic inflammation, termed inflamm-aging (*9*), the common association of bladder disorders with both aging and chronic inflammation suggests that an underlying driver of pathology may be age-associated inflammation (*10, 11*).

The bladder is a storage organ with a mucosal barrier that provides protection from both urinary wastes and pathogens (*12, 13*). In contrast to the epithelial barriers of other mucosal tissues, such as the intestinal epithelium, the multi-layered bladder epithelium (known as the urothelium) must be watertight and highly impermeable to solutes, metabolites, and toxic wastes present in the urine (*13*). Large superficial facet cells, the outer layer of the urothelium, form this barrier via synthesis of surface uroplakins and tight junctions that limit exposure of the underlying cells to urine. These superficial cells also have cell-autonomous defenses such as exfoliation and rapid regeneration in response to injury or infection (*12, 14*). Other epithelial barriers, including the intestine, lung, and eye, exhibit increased permeability with aging that is thought to stimulate age-associated inflammation (*15, 16*). In the bladder, disruption or dysfunction of the urothelial barrier leads to chronic inflammation and disorders such as IC/BPS and rUTIs (*7, 14, 17*). Whether urothelial barrier function is affected by aging is not known.

In contrast to other mucosal tissues, the bladder contains a relatively sparse repertoire of resident and patrolling immune cells (*12, 18*). During homeostasis, bladder immune cells consist of ∼70% antigen-presenting macrophages and dendritic cells, ∼10% T cells, and smaller numbers of NK cells, mast cells, eosinophils, and patrolling monocytes (*19, 20*). The bladder lacks dedicated mucosal secondary lymphoid organs (SLOs) that form during development like the Peyer’s patches of the small intestine. However, non-lymphoid tissues may form ectopic SLO-like structures, known as tertiary lymphoid tissues (TLTs), in response to chronic inflammation and antigen exposure (*21, 22*). Aging affects the size, presence, and functionality of TLTs in several tissues (*23-26*). For example, isolated lymphoid follicles (intestinal TLT) in aged mice have altered cellular compositions and produce more IgA compared to young mice (*27*), and inducible bronchus-associated lymphoid tissue (lung TLT) forms more robustly in response to cigarette smoke in aged mice compared to young mice (*28*). Whether aging affects immunity in the bladder mucosa is not known. Since both infectious and non-infectious chronic bladder inflammation is highly prevalent in older women, age-associated disruption of immune homeostasis in the bladder may mediate inflammatory pathology and lower urinary tract symptoms.

Here, we report that bladders from aged mice exhibit transcriptional signatures enriched in immune-mediated responses and a cellular repertoire skewed towards an expansion of lymphoid populations relative to bladders from young mice. Furthermore, we identify lymphoid cells organized into TLTs that we term bladder tertiary lymphoid tissues (bTLTs). bTLTS are found predominantly in females and analogous bTLTs are found in bladders of older women, many of whom had a history of rUTI. While aged mice similarly have a higher frequency of UTI recurrences than young mice, we demonstrate that bTLT form in an age-dependent manner that is independent of infection. We further demonstrate that bTLTs are capable of producing IgA^+^ plasma cells that form within germinal centers and secrete IgA into the urine. Moreover, we identify that TNFα is a major driver of bTLT formation, as TNFα-deficient mice lack bTLTs at any age. Finally, both aged wild type (WT) and TNFα^-/-^ mice have increased urothelial permeability. Thus, age-dependent TNFα responses to urothelial barrier dysfunction may ultimately drive chronic inflammation resulting in organized bTLTs in elderly women.

## RESULTS

### Adaptive immune networks distinguish bladders of aged mice from those of young mice

Since chronic, age-related bladder inflammation primarily affects women, we assessed how aging affects the global environment of the female bladder by performing RNA-sequencing on bladder tissue from young (3-4 month [mo]) and aged (18-22 mo) female mice. Compared to the transcriptome of bladders from young mice, bladders from aged mice had at least a 2-fold (FDR-adjusted P<0.05) increase in expression of 417 genes and decrease in expression of 59 genes (**Fig. 1A**). We then performed gene set enrichment analysis using the KEGG pathway database and identified 13 up-regulated pathways and 1 down-regulated pathway enriched in the bladder transcriptome of aged mice compared to those of young mice (**Fig. 1B**). Notably, the up-regulated pathways included B- and T-cell receptor signaling, antigen presentation, and IgA production pathways (**Fig. 1B**), which strongly suggested that the predominant age-associated changes to the bladder were immune-mediated responses.

**Fig. 1.**
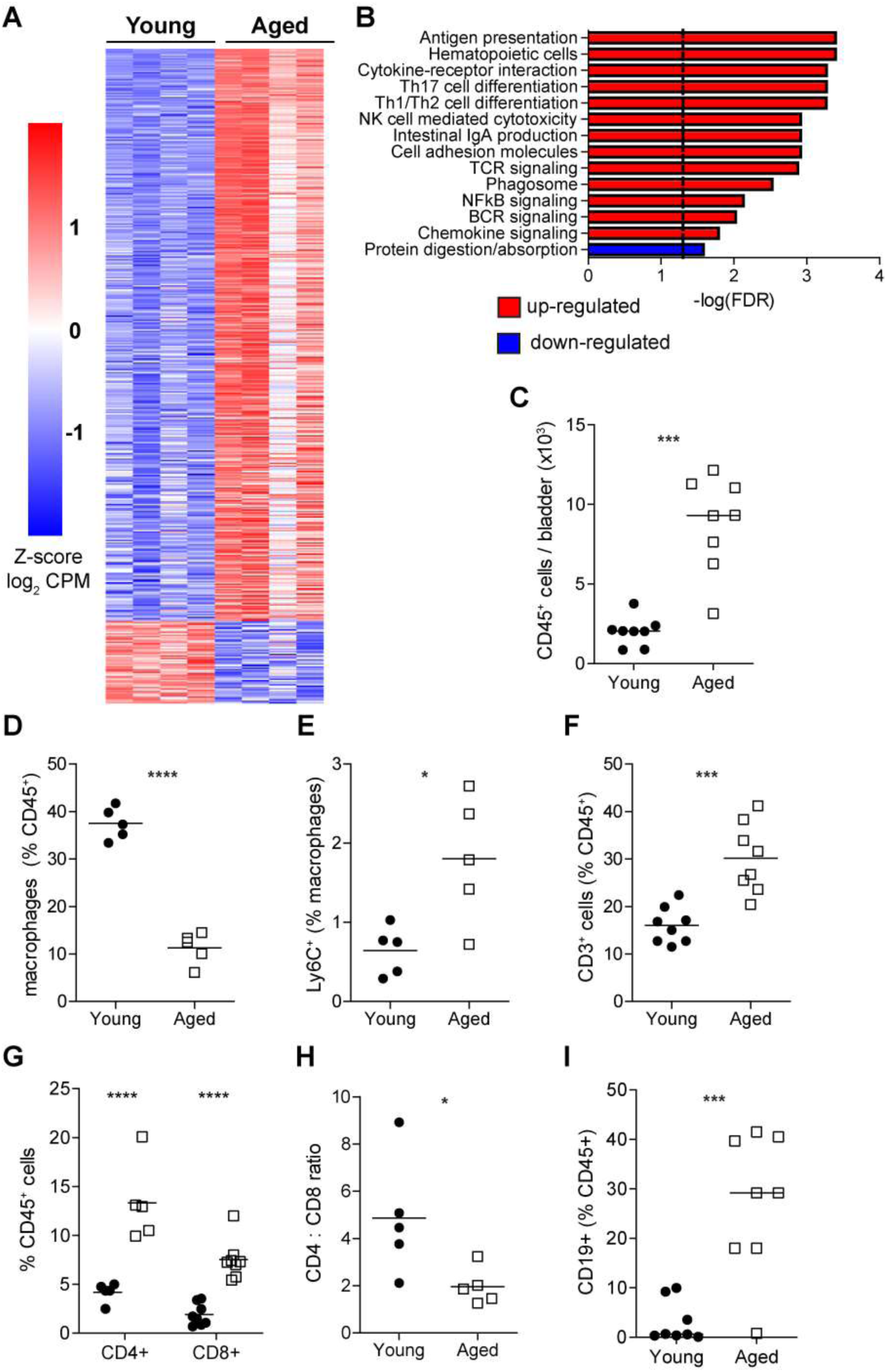
Immune processes and lymphocyte populations are expanded in bladders from aged mice. (**A**) Relative gene expression of whole bladder from young (3 mo, n=4) and aged mice (22 mo, n=4) showing genes with at least a 2-fold change (FDR-adjusted p<0.05). (**B**) Gene set enrichment analysis of KEGG pathways with FDR-adjusted p-values for up-(red) or down-regulated (blue) pathways. (**C**) Total number of live CD45^+^ cells in bladders (n=7 young, n=8 aged). Data are combined from 3 independent experiments. (**D**) Frequency of F4/80^+^CD64^+^ macrophages among total CD45^+^ cells in bladders (n=5/group). Data are combined from 2 independent experiments. (**E**) Frequency of Ly6C^+^ macrophages from bladders in (D). (**F**) Frequency of CD3^+^ T cells in bladders (n=8/group). Data are combined from 3 independent experiments. (**G**) Frequency of CD4^+^ and CD8^+^ T cells in bladders (n=5/group for CD4^+^ cells, n=8/group for CD8^+^ cells). Data are combined from 2 independent experiments. (**H**) Ratio of CD4^+^ T cell frequency to CD8^+^ T cell frequency in bladders (n=5/group). Data are combined from 2 independent experiments. (**I**) Frequency of CD19^+^ B cells in bladders (n=8/group). Data are combined from 3 independent experiments. ****p<0.0001, ***p<0.001, **p<0.01, *p<0.05 by Mann-Whitney U-test shown with median (C, H, I), unpaired t-test shown with mean (D, E), unpaired t-test of log-transformed data shown with geometric mean (F), or two-way ANOVA with Bonferroni post-test (G).

To determine if there were corresponding changes on a cellular level, we examined the immune compartment of bladders from young and aged mice by flow cytometry. Aged mice had higher numbers of immune cells in their bladders than young mice (**Fig. 1C**). Since macrophages are usually the largest immune cell population in the bladder and act as key sentinels that respond to infection and injury (*19, 20, 29, 30*), we examined whether bladder macrophages in aged mice differ from those in young mice. Despite the overall increase in immune cell numbers, there was a surprising decrease in the frequency of F4/80^+^CD64^+^ macrophages in bladders from aged mice compared to those from young mice (**Fig. 1D**). While the majority of bladder macrophages from both young and aged mice were the resident, Ly6C^-^ type, bladders from aged mice had a higher frequency of Ly6C^+^ macrophages, which are inflammatory monocyte-derived macrophages recently infiltrated into the tissue (**Fig. 1E**) (*19, 20, 31*). Since macrophages were no longer the predominant immune cell within the bladders of aged mice, we searched for expanded numbers of other immune cell types. Examining the lymphoid compartment, we found that bladders from aged mice were comprised of higher frequencies of T cells compared to those from young mice (**Fig. 1F**). Among both total immune cells and total T cells, bladders from aged mice had higher frequencies of CD4^+^ and CD8^+^ T cells than young mice (**Fig. 1G, S1**). Furthermore, the ratio of CD4^+^ T cells to CD8^+^ T cells was lower in aged mice than in young mice (**Fig. 1H**); this shift towards increased numbers of CD8^+^ T cells is a characteristic feature of age-associated inflammation, suggesting that the bladder mucosa may be affected by inflamm-aging (*32*). Whereas B cells are essentially absent in bladders from young mice, there was a substantial number of CD19^+^ B cells in bladders from aged mice (**Fig. 1I**). Thus, in aged mice, the expansion of lymphocytes displaces macrophages as the predominant immune cell within bladder tissue and may be a sign of age-associated inflammation in the bladder.

### Lymphocytes organize into tertiary lymphoid tissues in bladders of aged female mice and older women

Since bladder tissue differs between sexes, and females are known to develop more pronounced inflammatory phenotypes with age, we examined bladder tissue from young and aged mice of both sexes to localize the expanded lymphoid compartment observed by flow cytometry. Surprisingly, lymphoid cells in bladders from aged female mice were concentrated in large, dense aggregates throughout the bladder (**Fig. 2A**); in contrast, bladders from aged male mice rarely contained these structures and did not exhibit signs of tissue inflammation (**Fig. S2**). These findings mirror the epidemiology of chronic bladder inflammation; OAB, IC/BPS, and rUTIs are highly prevalent among older women and rarely found in older men. Since bladders from aged female mice, but not those from aged male mice, exhibited a dramatic inflammatory phenotype, we further studied age-associated inflammation in the female bladder as a model of what is seen in human populations.

**Fig. 2.**
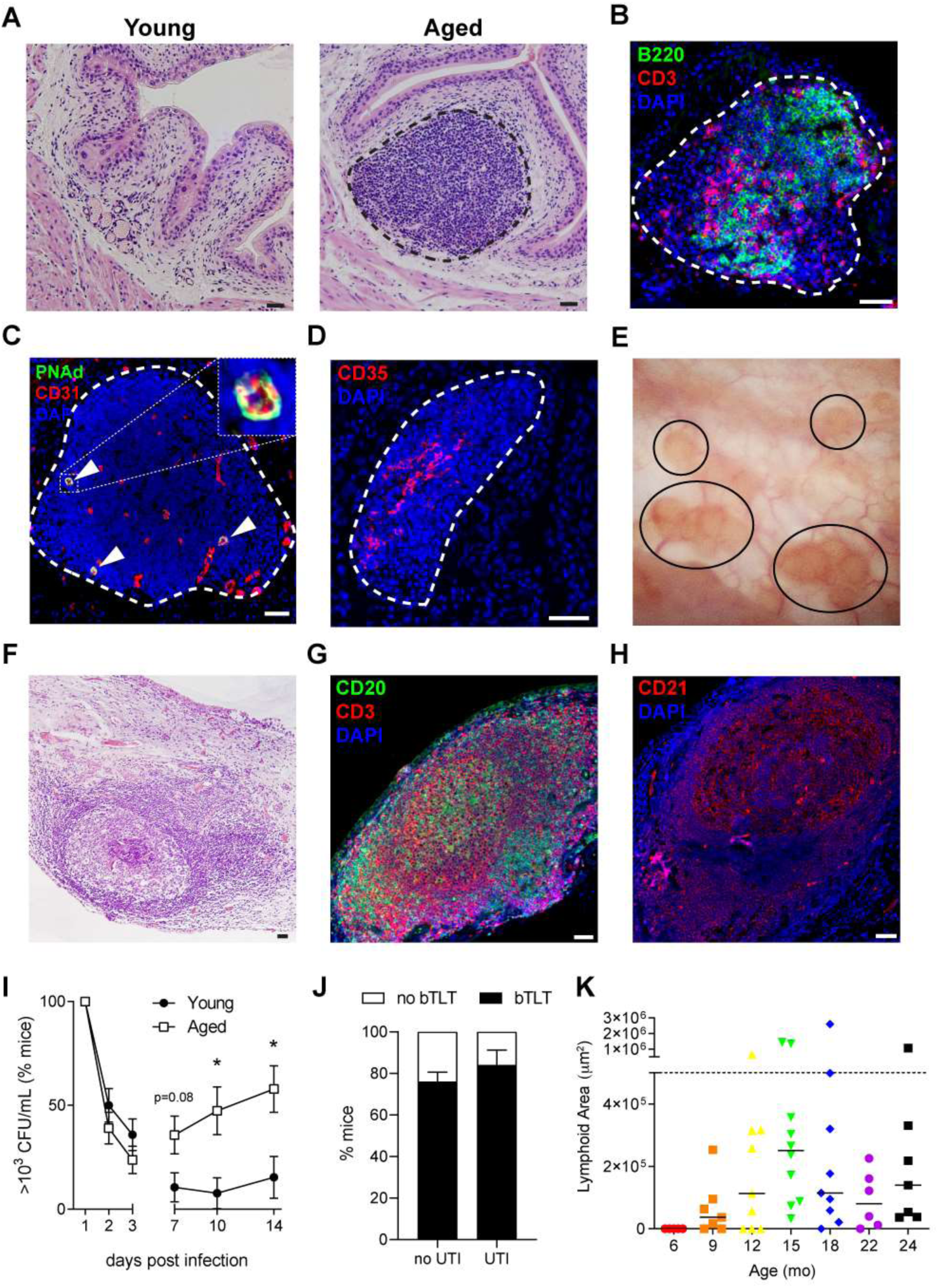
Bladder tertiary lymphoid tissue (bTLT) is found in aged mice and older women with a history of recurrent urinary tract infection. (**A**) Hematoxylin and eosin (H&E) image of bladder tissue from young (3 mo) and aged (18 mo) mice with an example of well-formed bTLT in aged mice. (**B**) Immunofluorescence (IF) image of B cells (B220, green) and T cells (CD3, red) within bTLT of aged mice. (**C**) IF image of high endothelial venules marked by peripheral node addressin (PNAd, green) and CD31 (red). (**D**) IF image of follicular dendritic cell (FDC) network marked by CD35 (red). (**E**) Cystoscopic image of nodules (black circles) in a chronically inflamed bladder that was biopsied. (**F**) H&E image of a well-formed lymphoid follicle in a bladder biopsy. (**G**) IF image of B cells (CD20, green) and T cells (CD3, red) in a bladder biopsy. (**H**) IF image of FDC network marked by CD21 (red) in a bladder biopsy. (**I**) Proportion of mice with urine titer >10^3^ CFU/mL uropathogenic *E. coli* at given time points (n=19-42 mice/group/time point). (**J**) Proportion of mice with bTLT before (no UTI, n=88) or after (UTI, n=25) infection. (**K**) Total area of bTLTs from mice of different ages (n=5-10 mice/age). Mouse images are representative of at least 5 mice. Human images are representative of n=12 patients. Dashed lines encircle bTLTs. All nuclei are stained blue with DAPI. Scale = 50 µm. *p<0.05, **p<0.01 by Fisher’s exact test shown with percentage and SEM (I) or Kruskal-Wallis with Dunn’s multiple comparison test shown with median.

The morphologic feature of the lymphoid infiltrates found in aged female bladders and the altered cell populations observed by flow cytometry (**Fig. 1F-I**) suggested that bladders from aged mice may contain tertiary lymphoid tissues (TLTs), which resemble the composition and structure of the secondary lymphoid organs (SLOs), but form ectopically in chronically inflamed tissues (*21, 22*). Nearly all of the B and T cells were localized to the lymphoid aggregates by immunofluorescence (**Fig. 2B**). B and T cells were segregated into distinct zones, forming an organized structure characteristic of TLTs. Other structural features of TLTs that are otherwise only found in dedicated lymphoid tissues include specialized high endothelial venules (HEVs), which permit the extravasation of migrating naive lymphocytes, and follicular dendritic cell (FDC) networks, which support the formation and maintenance of the B cell follicle by secretion of chemokines and capture of antibody via complement receptors (*33, 34*). We identified both HEVs, marked by co-expression of platelet endothelial cell adhesion molecule (PECAM; CD31^+^) and peripheral node addressin (PNAd^+^), and FDC networks, marked by complement receptor 1 (CR1; CD35^hi^), within large lymphoid aggregates (**Fig. 2C-D**), indicating that bladders from aged female mice contained *bone fide* TLTs. Together, our data demonstrate that B and T cells in bladders from aged mice localize to distinct aggregates with the organization and specialized structures characteristic of TLTs. Hereafter, these structures are termed bladder tertiary lymphoid tissues (bTLTs).

Small clinical case series occasionally report lymphoid aggregates or follicles in the bladder (*35-37*), but their relation to aging and outcome of chronic bladder inflammation remains unknown. Small, raised, reddish-yellow nodules can be grossly visualized by bladder endoscopy (cystoscopy) in some women with chronic bladder inflammation (**Fig. 2E**) (*39*); however, the significance of these nodules is not clear. Since aging is associated with high rates of chronic bladder inflammation in women, we sought to determine if bTLTs could be found in symptomatic women undergoing cystoscopy with possible biopsy. Since chronic bladder inflammation is most prevalent in older women, we hypothesized that these nodules are similar to bTLTs found in aged mice. Thirteen women with nodular lesions visualized by cystoscopy (**Fig. 2E**) were biopsied for pathologic diagnosis. Eleven of the 13 biopsies were read by a clinical pathologist and reported to have chronic inflammation with no malignant or pre-malignant changes. Furthermore, 7 of the 11 (64%) biopsies were noted in pathology reports to contain distinct lymphoid follicles in the lamina propria (**Fig. 2F**). Since some biopsies were too small to identify distinct lymphoid follicles by routine histology, we further analyzed separate biopsies obtained simultaneously for research. In biopsies from 12 of the 13 patients (92%), we identified distinct B and T cell organization (**Fig. 2G**) and FDC networks within follicles (**Fig. 2H**) similar to bTLTs found in aged mice. Since biopsies of healthy bladder tissue or bladders without visible lesions are not clinically indicated, it is impossible to determine if aging alone plays a role in the formation or enlargement of bTLTs from our cohort; however, it is notable that the median age of the patients with bladder biopsies was 64 years old (range 27-87). Women could develop bTLTs as a consequence of aging or may be predisposed to developing grossly-visible bTLTs in response to chronic bladder inflammation caused by rUTIs, IC/BPS, or OAB. Considering the striking similarity of bTLTs found in older women and aged mice, they may share underlying molecular drivers that could be further elucidated in this mouse model.

### Aged mice are susceptible to recurrent UTIs, but bTLTs form independent of infection

Ten out of the 13 women with bladder biopsies reported a history of recurrent UTIs (rUTIs), defined as at least 2 culture-positive UTIs in the past 6 months or 3 culture-positive UTIs in the past year. However, since biopsies are taken when there is not an acute infection, bTLTs in these patients represent a long-term inflammatory response in the bladder. In mice, chronic bladder inflammation can result in long-term changes to the mucosa that predisposes to further infections (*38, 40, 41*); thus we hypothesized that age-associated bTLTs may be associated with increased susceptibility to rUTIs. To determine if aged mice with bTLTs were similarly susceptible to rUTIs, we infected young and aged mice with uropathogenic *E. coli* and monitored urine bacterial titers over 14 days post infection (dpi). Young and aged mice were equally infected, as determined by urine bacterial titers at 6 hours post infection (**Fig. S3**). Over the first 3 dpi, young and aged mice cleared UPEC from their urine at equal rates; however, similar to older women, aged mice had more spontaneous recurrences of bacteriuria than young mice between 7 and 14 dpi (**Fig. 2I**). Histologically examining the bladders from aged mice, we found no difference in the proportion of mice that had bTLTs before and after infection (84% uninfected vs. 76% infected, p=0.5854, **Fig. 2J**). These findings suggested that, while aged mice are more susceptible to recurrent UTIs than young mice, bTLTs likely form due to age-associated inflammation rather than infection-induced inflammation. To determine when during the lifespan bTLT form in mice, we examined bladders from mice ranging from 3 to 24 mo of age. While no bTLTs (well-defined aggregates ≥10^4^ µm^2^) were found in mice 6 mo or younger, bTLTs were found in bladders beginning at 9 mo, and nearly all bladders contained bTLTs by 15 mo (**Fig. 2K**). While the number of bTLTs found in the bladder increased over time, the average size of each individual bTLT not significantly (**Fig. S3**). Thus, in mice, bTLTs form during normal aging beginning around 9 mo of age and increase in number, but not size, over time.

### bTLTs contain germinal centers that support local B cell maturation and IgA production

In SLOs, the homeostatic lymphoid chemokines CXCL12, CXCL13, CCL19, and CCL21 attract naïve lymphocytes and organize their characteristic follicular structure (*34, 42, 43*). Ectopic expression of one or more of these chemokines is sufficient to induce TLTs in a permissive tissue environment (*44, 45*). Given that bTLTs are found in bladders from aged mice, we anticipated that one or more of the homeostatic chemokines may be acting in the bladder. Indeed, *Cxcl13* was identified by RNA-seq as one of the most highly upregulated genes (17.8-fold change, FDR-adjusted P=0.0125) in bladders from aged mice. Consistent with TLTs found in other mucosal tissues, both *Cxcl13* (13.2-fold change) and *Ccl19* (10-fold change) were both highly upregulated in bladders from aged mice compared to those of young mice, as measured by qRT-PCR (**Fig. 3A)**. *Cxcl12* and *Ccl21* expression were not higher in bladders from aged mice (**Fig. S4**). Since these chemokines use overlapping receptors and have some functional redundancies, it is possible that CXCL13 and CCL19 are sufficient to recruit and organize bTLTs in aged mice while CXCL12 and CCL21 are dispensable (*42, 43*).

**Fig. 3.**
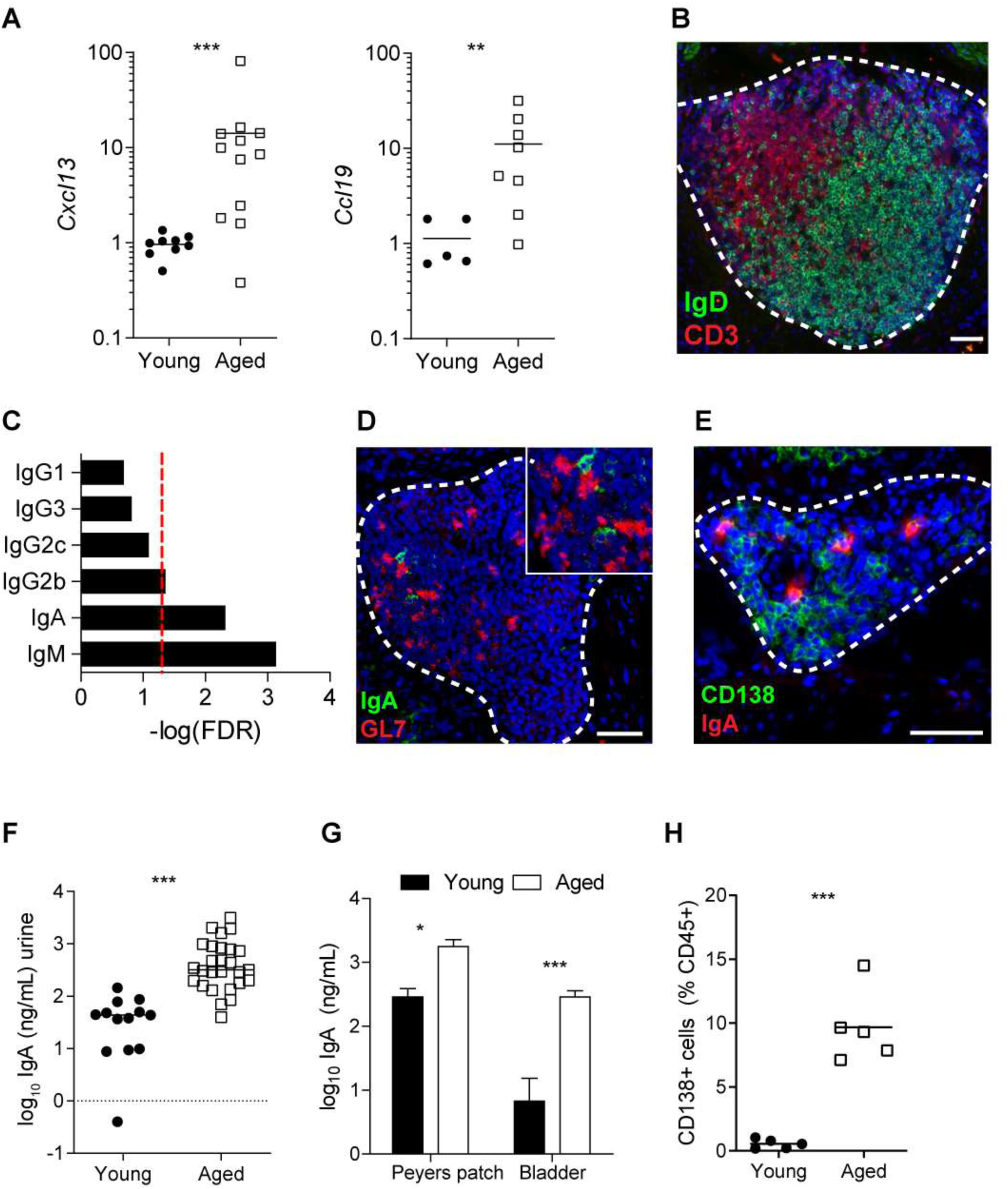
bTLTs in aged bladders generate local B cell responses. (**A**) Bladder gene expression of *Cxcl13* and *Ccl19* relative to young mice (n=6 young, n=12 aged). (**B**) Immunofluorescence (IF) image of T cells (CD3, red) and naive B cells (IgD, green) within bTLT from aged mice. (C) Enrichment of immunoglobulin (Ig) heavy chain constant genes in bladders from aged mice compared to those from young mice (n=4/group) in RNA-sequencing analysis. Red dashed line marks FDR-adjusted p=0.05. (**D**) IF image of bTLT with germinal center (GL7, red) containing IgA+ (green) cells. (**E**)) IF image of plasma cells (CD138, green) within bTLT. **(F)** Urine IgA concentration (n=13 young, n=27 aged). (**G**) IgA concentration after 24 hours *ex vivo* organ culture (n=5/group). Data are combined from 3 independent experiments. (**H**) Frequency of CD138^+^ plasma cells among total CD45^+^ cells in bladders (n=5/group). Data are combined from 2 independent experiments. All images are representative of at least 5 aged mice. Dashed lines encircle bTLTs. All nuclei are stained blue with DAPI. All scale bars are 50 µm. ****p<0.0001, ***p<0.001, **p<0.01, *p<0.05 by Mann-Whitney U-test shown with median (A, F, H) or 2-way ANOVA with Bonferroni post-test (G).

Since the homeostatic chemokines recruit naïve B and T cells to SLOs and TLTs, we hypothesized that bTLTs recruited naïve lymphocytes and generated *in situ* adaptive immune responses (*46, 47*). The majority of B cells within bTLTs stained positive for the naïve B cell marker IgD (**Fig. 3B**). After activation via antigen recognition, B cells may form germinal centers (GCs) within follicles, where they undergo somatic hypermutation, affinity maturation, and class switch recombination (*48*). If this were the case in the bladder, we would anticipate evidence of multiple isotypes of immunoglobulin (Ig) heavy chains. Indeed, RNA-seq analysis of bladders from young and aged mice revealed that the most highly upregulated genes in aged bladders were Ig constant genes, with IgM and IgA being the most significantly-upregulated (**Fig. 3C**). Since IgA is a class-switched isotype that is often produced in mucosal lymphoid tissues, we hypothesized that bTLTs supported local GCs that produce IgA-secreting plasma cells. Locally active GCs were identified within bTLTs by the highly-specific GC marker GL-7 (**Fig. 3D**), and CD138^+^IgA^+^ plasma cells were found localized to the edges of bTLTs (**Fig. 3E**). Furthermore, aged mice had 10-fold higher urine IgA concentrations than young mice (**Fig. 3F**), suggesting a role for bTLT in urinary IgA production. To determine if the increase in urine IgA was locally produced in the bladder, we cultured bladders from young and aged mice *ex vivo*. Bladders from aged mice secreted over 45 ng/mL more IgA than bladders from young mice (**Fig. 3G**), indicating that the increase in urine IgA was likely due to increased local production and secretion. The frequency of plasma cells in bladders from aged mice was also higher than those from young mice by flow cytometry (**Fig. 3H**), and IgA^+^ plasma cells were primarily localized to bTLT (**Fig. 3E**). These data further support a role for bTLTs in aged mice play in the local antibody responses and IgA production that is transported across the urothelium into the urine.

### Aging impairs urothelial barrier function and requires TNFα for bTLT formation

Both animal and human studies demonstrate an age-dependent loss of epithelial integrity in the gastrointestinal tract, lung, eye, and skin (*15*). Given the urothelium’s distinct role as a water- and solute-tight barrier to urinary wastes (*13, 49*), we hypothesized that aging impairs urothelial barrier integrity, which could stimulate chronic inflammation and bTLT formation in bladders from aged mice and older women. To test this hypothesis, we transurethrally instilled FITC-dextran into the bladders of young and aged mice (*50*). In bladders from both young and aged mice, FITC-dextran accumulated in the outermost, superficial urothelial cells (**Fig. 4A, white arrowheads**) that exclude urinary contents from underlying urothelial cells and tissue. However, FITC-dextran also penetrated into the intermediate and basal urothelial layers (**Fig. 4A, red arrows**) in aged mice but not in young mice Furthermore, fluorescence from FITC-dextran that had been absorbed through the urothelial basement membrane was higher in aged mice than in young mice (**Fig. 4B**). Thus, aged mice have increased penetration of urinary contents into the underlying bladder tissue, which could damage host proteins to stimulate bladder inflammation in an age-dependent manner (*49*).

**Fig. 4.**
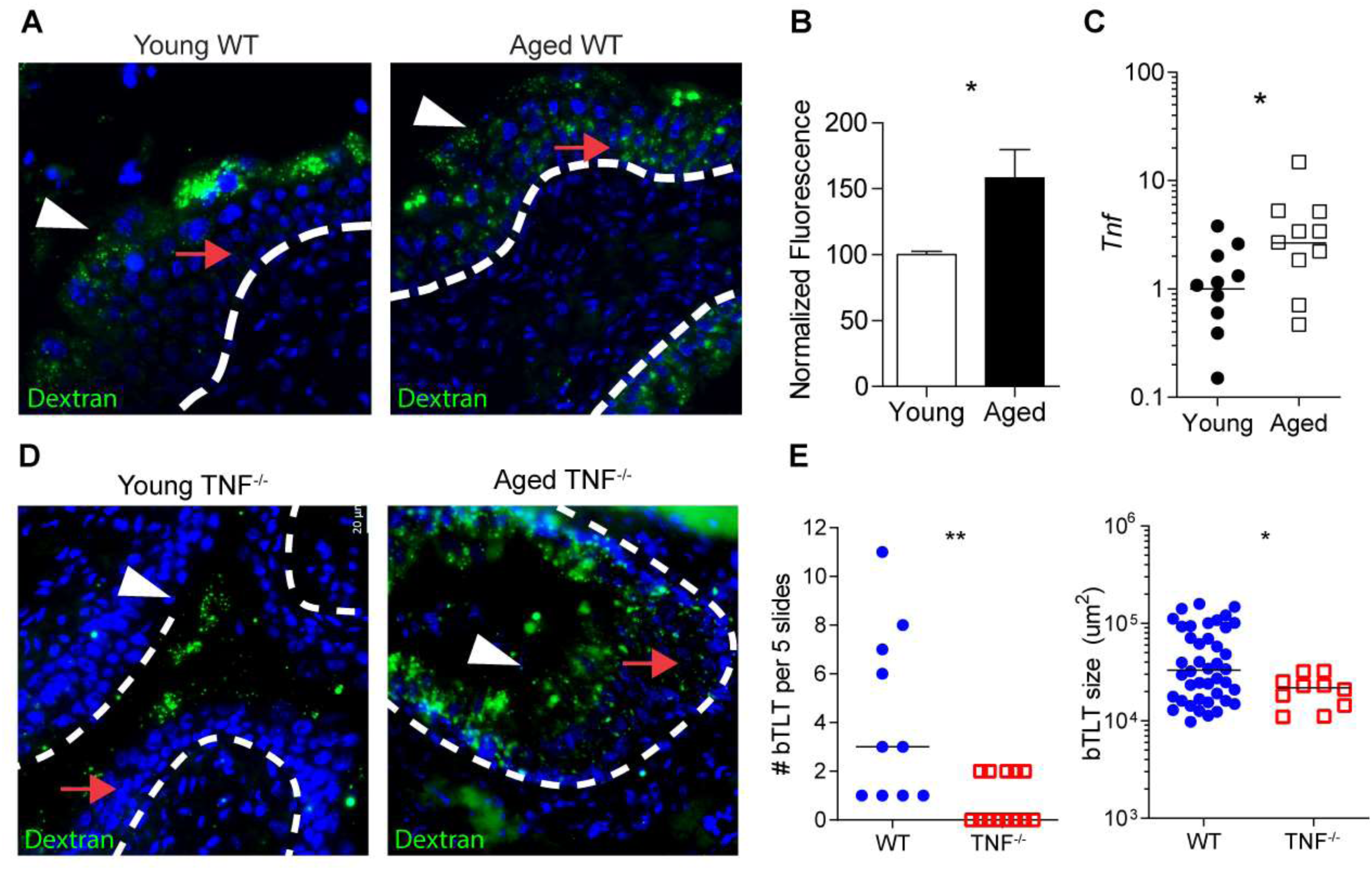
Aged TNFα^-/-^ mice lack bTLT but retain age-associated urothelial barrier defects. (**A**) FITC-dextran permeability in urothelium of young and aged mice. White arrowheads identify superficial umbrella cells with intracellular FITC-dextran. Red arrows identify basal and intermediate cell layers. (**B**) Mean gray value of stroma in mouse bladders treated with FITC-dextran (n=4/group). Values are normalized to the average value from young mice. Data are combined from 2 independent experiments. (**C**) Relative expression of *Tnf* in mouse bladder tissue (n=10/group)). (**D**) FITC-dextran permeability in urothelium of TNFα^-/-^ mice (n=2 young, n=3 aged) as in (A). (**E**) Number (left) and size (right) of bTLT in 5 bladder sections (n=10 WT, n=11 TNFα^-/-^). Images are representative of 2 independent experiments. All nuclei are stained blue with DAPI **p<0.01, *p<0.5 by Mann-Whitney U-test shown with median (B, E) or unpaired t-test of log-transformed data shown with geometric mean (C).

TNFα is a major pro-inflammatory cytokine involved in promoting inflamm-aging and its pathological consequences (*16, 51*). RNA-seq data indicated that *Tnf* was a top, locally-upregulated cytokine in bladders from aged mice (7.2-fold change, FDR-adjusted P=0.0035), which we confirmed by qRT-PCR (**Fig. 4C**). In the colon, age-associated TNFα has been shown to lead to epithelial permeability that permits increased translocation of bacterial products into the circulation, stimulating further age-associated inflammation via a positive feedback loop (*16, 52*). To test whether TNFα induces age-associated urothelial permeability, we transurethrally instilled FITC-dextran into the bladders of young and aged WT and TNFα^-/-^ mice. Aged TNFα^-/-^ had urothelial barrier defects similar to aged WT mice, thus TNFα was not required for age-associated urothelial permeability (**Fig. 4D**). However, bladders from aged TNFα^-/-^ mice rarely contained bTLT compared to age-matched WT mice (**Fig. 4E**), and these bTLT were significantly smaller than those found in WT mice (**Fig. 4F**). In contrast to the increased TNFα-dependent permeability in the aging gut (*16*), these data demonstrate that in the bladder, urothelial permeability increases in an age-dependent manner, while bTLT formation requires TNFα-dependent inflammatory responses.

## DISCUSSION

Postmenopausal and elderly women have increased susceptibility to bladder disorders involving chronic inflammation, including overactive bladder (OAB), interstitial cystitis/bladder pain syndrome (IC/BPS), and recurrent UTIs (rUTIs) (*1, 3, 4*); however, the underlying mechanisms predisposing older women to bladder inflammation have remained unclear (*10*). Here, we show the first characterization of age-induced changes to the immune system in the urinary bladder. Most strikingly, bladders from aged female mice, but not those from aged male mice, frequently contain large aggregates of lymphoid cells with a composition and organization consistent with tertiary lymphoid tissues (TLTs) (*21, 22, 33, 34*), thus we termed them bladder tertiary lymphoid tissues (bTLTs). We show that bTLTs develop with age in the absence of any experimental trigger, indicating that age itself is a risk factor for an increased inflammatory milieu in the bladder. bTLTs are not considered a normal finding in the bladder and are not found in healthy, young mice; thus bTLTs are a sign of chronic inflammation and perturbed homeostasis in this tissue (*36, 39, 53*). Furthermore, our finding that aging is also associated with urothelial barrier dysfunction and increased production of secretory IgA could shift the approach to studying and treating age-associated bladder inflammation. We also identified the key, age-associated inflammatory cytokine TNFα as a driver of bTLT formation in the aging bladder. These findings reveal a new target that could be used in the management of chronic bladder inflammation in older women. Furthermore, age-associated inflammation may promote bTLT formation and age-related susceptibility to chronic bladder inflammation.

Adaptive immune responses in lower urinary tract infections (e.g. cystitis without pyelonephritis) are reportedly limited, inadequate, and actively inhibited by the innate immune response to infection (*12, 19, 54*). Cellular characterization of aged bladders prior to infection revealed an influx of CD4^+^ and CD8^+^ T cells, naïve and activated B cells, and IgA^+^ plasma cells, all localized to organized bTLTs. Global bladder transcriptomes indicate that classical lymphoid neogenesis signaling pathways, including TNFα, lymphotoxin, and the homeostatic lymphoid chemokines, likely play a role in orchestrating the organization of these lymphocytes in aged bladders (*21, 22, 33, 34*). Young bladders do not contain substantial numbers of these cellular populations, particularly those of the B cell lineage (*12, 19*). T cell influx has been reported in young mice given multiple UTIs with uropathogenic *E. coli* (*55*), and aggregates of CD45^+^ cells have been observed in a subset of C3H/HeN mice with persistent bacteriuria, chronic cystitis, and pyelonephritis (*40*). Whether the adaptive immune cells that do infiltrate the bladder after infection remain resident in the tissue in the absence of on-going inflammation and antigenic stimulation hasn’t been addressed. Further studies will be needed to establish whether bladder lymphoid aggregates under these conditions are also organized bTLT that can support germinal center reactions and become a permanent feature of the bladder mucosae.

Since mucosal B cells and TLTs in other tissues can play both protective and pathogenic roles, it remains to be determined whether age-associated bTLTs are harmful or helpful (*21, 47, 56-60*). Here, we present evidence that bTLTs are active sites of naïve B cell recruitment, germinal center formation, and B cell maturation into plasma cells, suggesting that bTLTs serve as local antigen-processing centers in the bladder. Considering the association of bTLTs with aging and susceptibility to recurrent infection, we speculate that formation of age-associated bTLTs may exacerbate inflammatory pathology triggered by infection or other inflammatory insults in the bladder. While bTLTs have been observed in urothelial cancer specimens and correlated with advanced stages of disease (*61*), bladder cancer is less common in women than in men, and patients in our study had no evidence of malignancy or pre-malignant changes. TLTs associated with autoimmune diseases like rheumatoid arthritis typically propagate pathogenic auto-antibodies. Successful treatment with immune modulators such TNFα inhibitors can lead to TLT regression and reduction of pathology in rheumatoid arthritis patients (*56, 62, 63*). On the other hand, in other cancers, TLTs generally correlate with improved outcomes, implying that they aid in effective immune responses against tumor cells (*21, 65*). While adaptive immune responses from TLTs would intuitively be protective against infections, their association with chronic infections suggests that they are not always sufficient to eradicate such infections. In chronic hepatitis C virus infection, TLTs are associated with inflammatory pathology and autoimmune complications such as cryoglobulinemia (*64*). TLTs in the lung are associated with control and latency of *Mycobacterium tuberculosis* and thus thought to protect against reactivation of latent *M. tuberculosis* in this chronic infection (*66-68*). In chronic *Helicobacter pylori* gastroenteritis, TLTs have not been associated with specific infection outcomes, but these gastric TLTs have the potential to become mucosa-associated lymphoid tissue lymphomas (*69-71*). Interestingly, these gastric TLTs, both benign and malignant, frequently regress after eradication of *H. pylori* with antibiotics. The transient nature of *H. pylori*-associated gastric TLTs suggests that adaptive responses in this tissue may not be permanent. Thus, specific pathogens, tissue locations, and the nature of the immune responses occurring within TLTs impact whether TLTs are considered protective or pathogenic.

In the elderly, excessive inflammation in response to infection coupled with a lack of protective adaptive responses may indicate that TLTs associated with infection in this population could be pathogenic (*72*). TLTs in the elderly may also represent their inability to clear chronic infections and control latent infections (*73*). Given the high prevalence of UTIs among elderly women and the known mechanisms of uropathogenic *E. coli* persistence and recurrence, our discovery of age-associated bTLTs warrants further investigation from both a clinical and mechanistic perspective.

Mechanistically, we demonstrate that age-associated TNFα promotes the formation of bTLTs over the lifespan of mice. TNFα is a well-established marker of age-associated inflammation and likely has many roles in pathogenic changes that arise during older age (*11, 16*). Both aging and TNFα increase permeability in the intestinal epithelium (*16, 52*), but this phenomenon had not previously been demonstrated in the bladder epithelium. Given the critical importance of the bladder epithelium to serve as an impermeable barrier to urinary contents, increases in urothelial permeability have adverse consequences. For example, urothelial barrier defects are frequently found in interstitial cystitis/bladder pain syndrome (IC/BPS) patients, which is most often diagnosed in women over 40 (*3*). In an IC/BPS mouse model, urine exposure was required to induce permeability-mediated inflammation (*49*), and in another model, ectopic expression of TNFα in the bladder resulted in heighted pain sensitivity reminiscent of IC/BPS (*74*). In our studies, both aged WT and TNFα^-/-^ mice exhibited increased urothelial permeability, suggesting that age is the primary driver of epithelial dysfunction in the bladder. Since bladders from aged TNFα^-/-^ mice rarely contained bTLT, urothelial permeability is likely to be upstream of a TNFα-mediated inflammatory cascade. The cycle of age-induced epithelial permeability, urine exposure in the underlying tissue, and TNFα-mediated inflammatory responses may thus lead to or exacerbate bTLT formation. TNFα also plays a role in promoting the maturation of TLTs and germinal center reactions in SLOs (*75*), and could thus be a downstream factor limiting the formation of age-associated bTLTs. Further mechanistic studies are needed to fully define the exact role that TNFα plays in this process, which could lead to future improvement in the care and therapy of elderly women with chronic bladder inflammation.

As the global population ages, we must continue to assess how advanced age influences homeostasis and inflammatory responses. The newly recognized connection between aging, inflammation, epithelial permeability, and TLT formation could be a common theme affecting many mucosae and their age-related pathologies.

## Materials and Methods

### Mice

All experimental procedures were approved by the animal studies committee of Washington University in St. Louis School of Medicine (Animal Welfare Assurance #A-3381-01) and McMaster University’s Animal Research Ethics Board. 3- to 24-month-old C57B6/J mice were obtained from the National Institute of Aging. Mice were maintained under specified pathogen-free conditions in a barrier facility under a 12 h light-dark cycle. *Tnfa*^*-/-*^ and WT mice (originally from Jackson Laboratories) were bred and aged to 18-24 months at McMaster University (Theverajan 2017). To account for environmental factors, experiments with *Tnfa*^*-/-*^ mice were compared to WT mice raised in the same facility.

### Mouse Urinary Tract Infection

UTI89, a clinical UPEC isolate from a patient with recurrent cystitis was grown statically for 17 h in Luria-Bertani broth (Tryptone 10 g/L, Yeast extract. 5 g/L. and NaCl 10g/L) at 37°C prior to infection. Mice were anesthetized and inoculated via transurethral catheterization with 10^7^ colony forming units (CFUs) of UTI89 in phosphate-buffered saline (PBS; Sigma-Aldrich, D8537). Urines were collected at indicated time points and spotted onto LB-agar plates to measure bacterial titers.

### Histological and Immunofluorescence analysis

Bladders were aseptically removed, cut along the anterior-posterior axis, and fixed in 10% neutral buffered formalin or methacarn (60% methanol, 30% chloroform, 10% acetic acid), embedded in paraffin, stained with hematoxylin and eosin (7211, Richard-Allen Scientific) and imaged on a Nanozoomer 2.0-HT system (Hamamatsu). Matching anterior or posterior bladder halves were compared within groups. Number and area of bTLTs were determined in 5 sections spaced 150 µm apart using NDP,view2 software (Hamamatsu). Compact aggregates >10,000 µm^2^ were considered bTLTs. For immunofluorescence analysis, bladders were embedded in OCT Compound (4583, Tissue-Tek) and flash frozen. 7 µm sections were fixed with 1:1 methanol-acetone, rehydrated in PBS, and blocked with Avidin/Biotin Blocking Kit (SP2001, Vector Laboratories) followed by 1% BSA in PBS. Primary antibodies against B220 (13-0452, eBioscience), CD3 (14-0031-82, eBioscience) PNAd (120803, BioLegend), CD31 (ab28364, Abcam), CD138 (142511, BioLegend), CD35 (558768, BD Biosciences), GL7 (13-5902-81, eBioscience), and IgD (1120-01, SouthernBiotech) were incubated overnight at 4° and detected with streptavidin-conjugated and species-specific secondary antibodies followed by Hoechst dye. Slides were covered slipped with Prolong Gold antifade (P36930, Invitrogen) and imaged on a Zeiss Axio Imager M2 microscope with a Hamamatsu Flash4.0 camera using Zeiss Zen Pro software.

### Urothelial permeability assay

50 µL of 10 mg/mL 10 kD FITC-dextran (D1821, Invitrogen) was transurethrally inoculated into mice as previously described (Shin 2011). After 90 minutes, bladders were embedded in OCT Compound (4583, Tissue-Tek) and flash frozen. 7 µm sections were briefly dipped in 1:1 methanol-acetone and PBS, then cover-slipped with Prolong Diamond Antifade with DAPI (P36971, Invitrogen). Images were acquired using a fixed exposure set to detect fluorescence in young WT bladders. Stromal fluorescence was quantified by averaging the mean gray value for FITC channel in 6 random squares from 2 separate images for each mouse using ImageJ software. Values were combined for each mouse and used for statistical analysis.

### Organ culture and IgA ELISA

Bladders were aseptically removed, rinsed with PBS, bisected, and both halves cultured together in 500 µL RPMI-1640 with 10% FBS, Pen/Strep, 10 mM HEPES, and glutamax. Supernatants were removed after 24 hrs and cleared of debris by centrifugation. IgA concentration in urines and culture media was determined by ELISA according to manufacturer protocol (88-50450-22, Invitrogen).

### Flow cytometry

Bladders were aseptically removed, minced with scissors, and digested at 37° for 30 minutes in RPMI-1640 with 10mM HEPES, collagenase D (C5318, Sigma-Aldrich), and DNAse (10104159001, Sigma-Aldrich). Bladders were forced through a 70 µm cell strainer (352350, Corning) and washed with 5% FBS in PBS. Single cell suspensions were stained with anti-CD45-eFluor450 (48-0451-82, eBioscience), anti-CD3-APC (17-0032-82, eBioscience), anti-CD19-PE (115511, BioLegend), anti-CD4-FITC (100405, BioLegend), anti-CD8-PE/Cy7 (100721, BioLegend), anti-CD138-BrillantViolet605 (142515, BioLegend), and 7-AAD (420404, BioLegend). Data was acquired on LSR II flow cytometer (BD) and analyzed with FlowJo software v10.0. Gates were determined with isotype antibodies in bladder suspensions from young mice.

### RNA-sequencing

RNA was purified from snap frozen, homogenized bladders with RNeasy Mini Kit (74101, Qiagen) and RNase-free DNase digestion kit (79254, Qiagen). Libraries were prepared with Ribo-Zero rRNA depletion kit (Illumina) and sequenced on HiSeq3000 (Illumina). Reads were aligned to the Ensembl top-level assembly with STAR version 2.0.4b. Gene counts were derived from the number of uniquely aligned unambiguous reads by Subread:featureCount version 1.4.5. Transcript counts were produced by Sailfish version 0.6.3. Sequencing performance was assessed for total number of aligned reads, total number of uniquely aligned reads, genes and transcripts detected, ribosomal fraction, known junction saturation and read distribution over known gene models with RSeQC version 2.3. All gene-level and transcript counts were then imported into the R/Bioconductor package EdgeR and TMM normalization size factors were calculated to adjust samples for differences in library size. Ribosomal features as well as any feature not expressed in at least the smallest condition size minus one sample were excluded from further analysis and TMM size factors were recalculated to create effective TMM size factors. The TMM size factors and the matrix of counts were then imported into R/Bioconductor package Limma and weighted likelihoods based on the observed mean-variance relationship of every gene/transcript and sample were then calculated for all samples with the voomWithQualityWeights function. Performance of the samples was assessed with a spearman correlation matrix and multi-dimensional scaling plots. Gene/transcript performance was assessed with plots of residual standard deviation of every gene to their average log-count with a robustly fitted trend line of the residuals. Generalized linear models were then created to test for gene/transcript level differential expression. Differentially expressed genes and transcripts were then filtered for False Discovery Rate (FDR)-adjusted p-values less than or equal to 0.05. Pathways analysis was performed using results imported into the R/Bioconductor packages GAGE and Pathview.

### RT-qPCR

Bladders were flash frozen or stabilized in RNA Save (01-891-1A, Biological Industries) and RNA extracted using TRIzol reagent (15596018, Invitrogen) according to manufacturer protocol followed by gDNA digestion with TURBO DNA-free kit (AM1907, Invitrogen). cDNA was generated using Superscript III Reverse Transcriptase (18064014, Invitrogen). qPCR was performed with SsoAdvanced Universal SYBR Green Supermix (1725275, Bio-Rad) on a CFX96 Touch Real-Time PCR Detection System (Bio-Rad). Fold-changes were calculated using ΔΔCt method and normalized internally to 18S expression.

### Tissue analysis of human bladder biopsy samples

Cystoscopic pinch biopsies were obtained from patients undergoing gynecologic surgery with previous findings of bladder nodules from the Women’s Genitourinary Tract Specimen Consortium at Washington University in St. Louis School of Medicine (IRB#201810094) biorepository. Biopsies (approx. size 1 mm^3^) were fixed in 10% neutral buffered formalin, embedded in paraffin, stained with hematoxylin and eosin. Images were acquired with Olympus DP71 software. Sections were stained with antibodies to CD20 (14-0202-80), CD3 (ab5690, Abcam), and CD21 (NBP1-22527, Novus Biological) and imaged as above.

### Statistical analyses

Statistical tests were performed in GraphPad Prism 8. Data sets were evaluated for normality and lognormality with Anderson-Darling, D’Agostino-Pearson, Shipiro-Wilk, and Kolmogorov-Smirno tests. Lognormal distributions were log-transformed and analyzed as a parametric distribution. Unpaired t-tests (with Welch’s correction where appropriate) or two-way ANOVA with Bonferroni post-tests were used for parametric data and Mann-Whitney U test or Kruskal-Wallis with Dunn’s multiple comparison tests were used for non-parametric data. P<0.05 was considered significant. Data points represent individual animals. Lines represent the mean for normal distributions, geometric mean for log-normal distributions, or median for non-parametric distributions. Error bars represent SEM.

## Supporting information

Manuscript

## Acknowledgments

We thank Drs. Deborah Frank and Jason Mills for editorial comments; Drs. Melanie Meister, Stacy Lenger, and the Women’s Genitourinary Tract Specimen Consortium (WGUTSC) for biopsy procurement; and the Genome Technology Access Center (GTAC) for performing and processing sequencing data.

## Author contributions

Conceptualization MML, CW, IUM. Methodology MML, CW, IUM, DMEB. Investigation MML, CW, CS, END. Data curation ZJ. Data analysis MML. Resources JLL, DMEB. Supervision CW, DMEB, JLL, IUM. Funding IUM, DMEB. Visualization MML. Writing--original draft MML, IUM. Writing-review & editing MML, CW, DMEB, IUM

## Data and materials availability

All sequencing data will be deposited in an appropriate public repository.

## References

1. A. M. Suskind et al., Incidence and Management of Uncomplicated Recurrent Urinary Tract Infections in a National Sample of Women in the United States. Urology 90, 50–55 (2016).

2. J. A. Koziol, H. P. Adams, A. Frutos, Discrimination between the ulcerous and the nonulcerous forms of interstitial cystitis by noninvasive findings. The Journal of urology 155, 87–90 (1996).

3. L. J. Simon, J. R. Landis, D. R. Erickson, L. M. Nyberg, The Interstitial Cystitis Data Base Study: concepts and preliminary baseline descriptive statistics. Urology 49, 64–75 (1997).

4. I. Nygaard et al., Prevalence of symptomatic pelvic floor disorders in US women. JAMA 300, 1311–1316 (2008).

5. K. A. Kline, D. M. Bowdish, Infection in an aging population. Current opinion in microbiology 29, 63–67 (2016).

6. E. Ma et al., A multiplexed analysis approach identifies new association of inflammatory proteins in patients with overactive bladder. Am J Physiol Renal Physiol 311, F28–34 (2016).

7. L. Grundy, A. Caldwell, S. M. Brierley, Mechanisms Underlying Overactive Bladder and Interstitial Cystitis/Painful Bladder Syndrome. Front Neurosci 12, 931 (2018).

8. N. N. Maserejian et al., Incidence of lower urinary tract symptoms in a population-based study of men and women. Urology 82, 560–564 (2013).

9. C. Franceschi et al., Inflamm-aging. An evolutionary perspective on immunosenescence. Ann N Y Acad Sci 908, 244–254 (2000).

10. P. Tyagi et al., Association of inflammaging (inflammation + aging) with higher prevalence of OAB in elderly population. Int Urol Nephrol 46, 871–877 (2014).

11. C. Franceschi et al., Inflammaging 2018: An update and a model. Semin Immunol 40, 1–5 (2018).

12. S. N. Abraham, Y. Miao, The nature of immune responses to urinary tract infections. Nature reviews. Immunology 15, 655–663 (2015).

13. X. R. Wu, X. P. Kong, A. Pellicer, G. Kreibich, T. T. Sun, Uroplakins in urothelial biology, function, and disease. Kidney Int 75, 1153–1165 (2009).

14. I. U. Mysorekar, M. Isaacson-Schmid, J. N. Walker, J. C. Mills, S. J. Hultgren, Bone morphogenetic protein 4 signaling regulates epithelial renewal in the urinary tract in response to uropathogenic infection. Cell host & microbe 5, 463–475 (2009).

15. A. R. Parrish, The impact of aging on epithelial barriers. Tissue Barriers 5, e1343172 (2017).

16. N. Thevaranjan et al., Age-Associated Microbial Dysbiosis Promotes Intestinal Permeability, Systemic Inflammation, and Macrophage Dysfunction. Cell host & microbe 21, 455–466 e454 (2017).

17. R. E. Hurst et al., Increased bladder permeability in interstitial cystitis/painful bladder syndrome. Transl Androl Urol 4, 563–571 (2015).

18. M. A. Ingersoll, M. L. Albert, From infection to immunotherapy: host immune responses to bacteria at the bladder mucosa. Mucosal Immunol 6, 1041–1053 (2013).

19. G. Mora-Bau et al., Macrophages Subvert Adaptive Immunity to Urinary Tract Infection. PLoS pathogens 11, e1005044 (2015).

20. M. Schiwon et al., Crosstalk between sentinel and helper macrophages permits neutrophil migration into infected uroepithelium. Cell 156, 456–468 (2014).

21. C. Pitzalis, G. W. Jones, M. Bombardieri, S. A. Jones, Ectopic lymphoid-like structures in infection, cancer and autoimmunity. Nature reviews. Immunology 14, 447–462 (2014).

22. F. Aloisi, R. Pujol-Borrell, Lymphoid neogenesis in chronic inflammatory diseases. Nature reviews. Immunology 6, 205–217 (2006).

23. Y. Huang et al., Identification of novel genes associated with renal tertiary lymphoid organ formation in aging mice. PloS one 9, e91850 (2014).

24. Y. Sato et al., Heterogeneous fibroblasts underlie age-dependent tertiary lymphoid tissues in the kidney. JCI Insight 1, e87680 (2016).

25. P. Singh et al., Lymphoid neogenesis and immune infiltration in aged liver. Hepatology 47, 1680–1690 (2008).

26. R. Grabner et al., Lymphotoxin beta receptor signaling promotes tertiary lymphoid organogenesis in the aorta adventitia of aged ApoE-/-mice. J Exp Med 206, 233–248 (2009).

27. K. G. McDonald, M. R. Leach, C. Huang, C. Wang, R. D. Newberry, Aging impacts isolated lymphoid follicle development and function. Immun Ageing 8, 1 (2011).

28. G. John-Schuster et al., Inflammaging increases susceptibility to cigarette smoke-induced COPD. Oncotarget 7, 30068–30083 (2016).

29. J. W. Symington et al., ATG16L1 deficiency in macrophages drives clearance of uropathogenic E. coli in an IL-1beta-dependent manner. Mucosal Immunol 8, 1388–1399 (2015).

30. A. J. Carey et al., Uropathogenic Escherichia coli Engages CD14-Dependent Signaling to Enable Bladder-Macrophage-Dependent Control of Acute Urinary Tract Infection. J Infect Dis 213, 659–668 (2016).

31. A. Dixit et al., Frontline Science: Proliferation of Ly6C(+) monocytes during urinary tract infections is regulated by IL-6 trans-signaling. J Leukoc Biol 103, 13–22 (2018).

32. F. Zanni et al., Marked increase with age of type 1 cytokines within memory and effector/cytotoxic CD8+ T cells in humans: a contribution to understand the relationship between inflammation and immunosenescence. Exp Gerontol 38, 981–987 (2003).

33. G. W. Jones, D. G. Hill, S. A. Jones, Understanding Immune Cells in Tertiary Lymphoid Organ Development: It Is All Starting to Come Together. Frontiers in immunology 7, 401 (2016).

34. D. L. Drayton, S. Liao, R. H. Mounzer, N. H. Ruddle, Lymphoid organ development: from ontogeny to neogenesis. Nature immunology 7, 344–353 (2006).

35. S. Nekkanti, A. Doering, D. L. Zynger, A. F. Hundley, von Brunn’s Nests and Follicular Cystitis Following Intradetrusor OnabotulinumtoxinA Injections for Overactive Bladder. Urol Case Rep 14, 38–41 (2017).

36. T. A. Schlager, R. LeGallo, D. Innes, J. O. Hendley, C. A. Peters, B cell infiltration and lymphonodular hyperplasia in bladder submucosa of patients with persistent bacteriuria and recurrent urinary tract infections. The Journal of urology 186, 2359–2364 (2011).

37. F. P. Marsh, R. Banerjee, P. Panchamia, The relationship between urinary infection, cystoscopic appearance, and pathology of the bladder in man. J Clin Pathol 27, 297–307 (1974).

38. V. P. O’Brien et al., A mucosal imprint left by prior Escherichia coli bladder infection sensitizes to recurrent disease. Nat Microbiol 2, 16196 (2016).

39. C. Stirling, J. E. Ash, Chronic proliferative lesions of urinary tract. The Journal of urology 3, 342–360 (1941).

40. T. J. Hannan, I. U. Mysorekar, C. S. Hung, M. L. Isaacson-Schmid, S. J. Hultgren, Early severe inflammatory responses to uropathogenic E. coli predispose to chronic and recurrent urinary tract infection. PLoS pathogens 6, e1001042 (2010).

41. T. J. Hannan et al., Inhibition of Cyclooxygenase-2 Prevents Chronic and Recurrent Cystitis. EBioMedicine 1, 46–57 (2014).

42. G. Muller, P. Reiterer, U. E. Hopken, S. Golfier, M. Lipp, Role of homeostatic chemokine and sphingosine-1-phosphate receptors in the organization of lymphoid tissue. Ann N Y Acad Sci 987, 107–116 (2003).

43. S. A. Luther et al., Differing activities of homeostatic chemokines CCL19, CCL21, and CXCL12 in lymphocyte and dendritic cell recruitment and lymphoid neogenesis. Journal of immunology 169, 424–433 (2002).

44. S. C. Chen et al., Ectopic expression of the murine chemokines CCL21a and CCL21b induces the formation of lymph node-like structures in pancreas, but not skin, of transgenic mice. Journal of immunology 168, 1001–1008 (2002).

45. S. A. Luther, T. Lopez, W. Bai, D. Hanahan, J. G. Cyster, BLC expression in pancreatic islets causes B cell recruitment and lymphotoxin-dependent lymphoid neogenesis. Immunity 12, 471–481 (2000).

46. G. W. Jones, S. A. Jones, Ectopic lymphoid follicles: inducible centres for generating antigen-specific immune responses within tissues. Immunology 147, 141–151 (2016).

47. K. A. Knoop, R. D. Newberry, Isolated Lymphoid Follicles are Dynamic Reservoirs for the Induction of Intestinal IgA. Frontiers in immunology 3, 84 (2012).

48. L. Mesin, J. Ersching, G. D. Victora, Germinal Center B Cell Dynamics. Immunity 45, 471–482 (2016).

49. R. Soler et al., Urine is necessary to provoke bladder inflammation in protamine sulfate induced urothelial injury. The Journal of urology 180, 1527–1531 (2008).

50. K. Shin et al., Hedgehog/Wnt feedback supports regenerative proliferation of epithelial stem cells in bladder. Nature 472, 110–114 (2011).

51. C. Franceschi et al., Inflammaging and anti-inflammaging: a systemic perspective on aging and longevity emerged from studies in humans. Mech Ageing Dev 128, 92–105 (2007).

52. T. Y. Ma et al., TNF-alpha-induced increase in intestinal epithelial tight junction permeability requires NF-kappa B activation. Am J Physiol Gastrointest Liver Physiol 286, G367–376 (2004).

53. S. Hansson et al., Follicular cystitis in girls with untreated asymptomatic or covert bacteriuria. The Journal of urology 143, 330–332 (1990).

54. C. Y. Chan, A. L. St John, S. N. Abraham, Mast cell interleukin-10 drives localized tolerance in chronic bladder infection. Immunity 38, 349–359 (2013).

55. P. Thumbikat, C. Waltenbaugh, A. J. Schaeffer, D. J. Klumpp, Antigen-specific responses accelerate bacterial clearance in the bladder. Journal of immunology 176, 3080–3086 (2006).

56. J. Rangel-Moreno et al., Inducible bronchus-associated lymphoid tissue (iBALT) in patients with pulmonary complications of rheumatoid arthritis. J Clin Invest 116, 3183–3194 (2006).

57. T. Eddens et al., Pneumocystis-Driven Inducible Bronchus-Associated Lymphoid Tissue Formation Requires Th2 and Th17 Immunity. Cell Rep 18, 3078–3090 (2017).

58. R. G. Lorenz, R. D. Newberry, Isolated lymphoid follicles can function as sites for induction of mucosal immune responses. Ann N Y Acad Sci 1029, 44–57 (2004).

59. D. Lucchesi, M. Bombardieri, The role of viruses in autoreactive B cell activation within tertiary lymphoid structures in autoimmune diseases. J Leukoc Biol 94, 1191–1199 (2013).

60. K. Neyt, F. Perros, C. H. GeurtsvanKessel, H. Hammad, B. N. Lambrecht, Tertiary lymphoid organs in infection and autoimmunity. Trends Immunol 33, 297–305 (2012).

61. M. Koti et al., Tertiary Lymphoid Structures Associate with Tumour Stage in Urothelial Bladder Cancer. Bladder Cancer 3, 259–267 (2017).

62. J. H. Anolik et al., Cutting edge: anti-tumor necrosis factor therapy in rheumatoid arthritis inhibits memory B lymphocytes via effects on lymphoid germinal centers and follicular dendritic cell networks. Journal of immunology 180, 688–692 (2008).

63. J. D. Canete et al., Clinical significance of synovial lymphoid neogenesis and its reversal after anti-tumour necrosis factor alpha therapy in rheumatoid arthritis. Ann Rheum Dis 68, 751–756 (2009).

64. , (!!! INVALID CITATION !!! (Sansonno et al., 2008)).

65. H. J. Kim et al., Establishment of early lymphoid organ infrastructure in transplanted tumors mediated by local production of lymphotoxin alpha and in the combined absence of functional B and T cells. Journal of immunology 172, 4037–4047 (2004).

66. S. A. Khader et al., IL-23 is required for long-term control of Mycobacterium tuberculosis and B cell follicle formation in the infected lung. Journal of immunology 187, 5402–5407 (2011).

67. S. A. Khader et al., In a murine tuberculosis model, the absence of homeostatic chemokines delays granuloma formation and protective immunity. Journal of immunology 183, 8004–8014 (2009).

68. T. Ulrichs et al., Human tuberculous granulomas induce peripheral lymphoid follicle-like structures to orchestrate local host defence in the lung. J Pathol 204, 217–228 (2004).

69. M. Kobayashi et al., Induction of peripheral lymph node addressin in human gastric mucosa infected by Helicobacter pylori. Proceedings of the National Academy of Sciences of the United States of America 101, 17807–17812 (2004).

70. D. Sansonno et al., Increased serum levels of the chemokine CXCL13 and up-regulation of its gene expression are distinctive features of HCV-related cryoglobulinemia and correlate with active cutaneous vasculitis. Blood 112, 1620–1627 (2008).

71. N. Asano, K. Iijima, T. Koike, A. Imatani, T. Shimosegawa, Helicobacter pylori-negative gastric mucosa-associated lymphoid tissue lymphomas: A review. World journal of gastroenterology : WJG 21, 8014–8020 (2015).

72. M. N. Bouchlaka et al., Aging predisposes to acute inflammatory induced pathology after tumor immunotherapy. J Exp Med 210, 2223–2237 (2013).

73. C. Franceschi, J. Campisi, Chronic inflammation (inflammaging) and its potential contribution to age-associated diseases. J Gerontol A Biol Sci Med Sci 69 Suppl 1, S4–9 (2014).

74. W. Yang, T. J. Searl, R. Yaggie, A. J. Schaeffer, D. J. Klumpp, A MAPP Network study: overexpression of tumor necrosis factor-alpha in mouse urothelium mimics interstitial cystitis. Am J Physiol Renal Physiol 315, F36–F44 (2018).

75. Y. Wang, J. Wang, Y. Sun, Q. Wu, Y. X. Fu, Complementary effects of TNF and lymphotoxin on the formation of germinal center and follicular dendritic cells. Journal of immunology 166, 330–337 (2001).

