## Supplementary material for "Tertiary lymphoid tissue develops during normal aging in mice and humans": Manuscript

**Supplementary Materials**

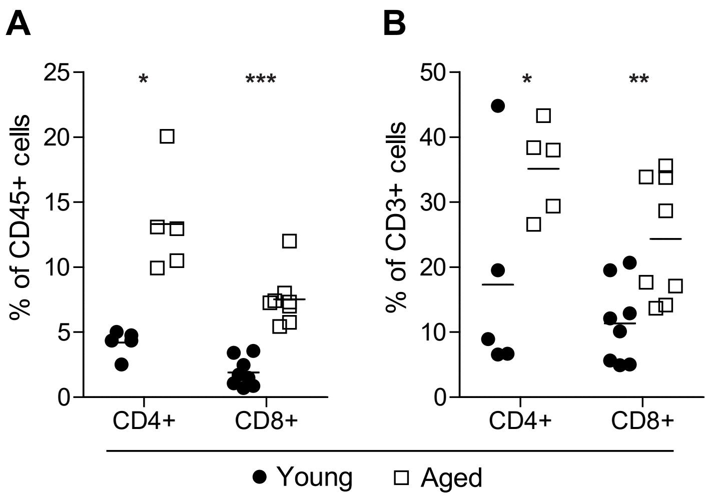


**Fig. S1 CD4+ and CD8+ T cells in young and aged mouse bladders.** (A) CD4+ and CD8+ subsets among total CD45+ cells. (B) CD4+ and CD8+ cells among all CD3+ cells. Data are combined from 2 independent experiments. n=5-8/group. *p<0.05, **p<0.01,

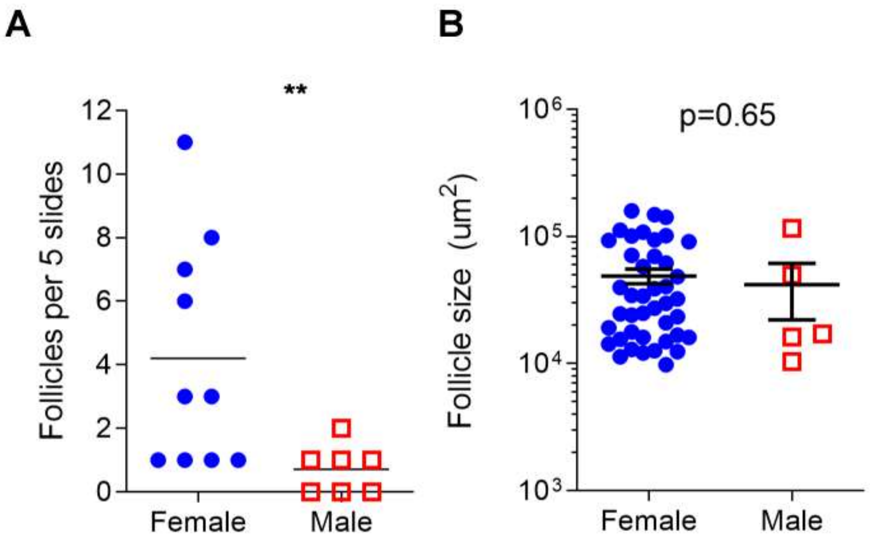


**Fig. S2. Male mice do not develop bTLT during aging. (A)** Number of bTLT (>$10^3$ $um^2$) in 20-22 mo old male and female mice. **(B)** Size of individual bTLT in 20-22 mo old male and female mice. **P<0.01 by Mann-Whitney U test.

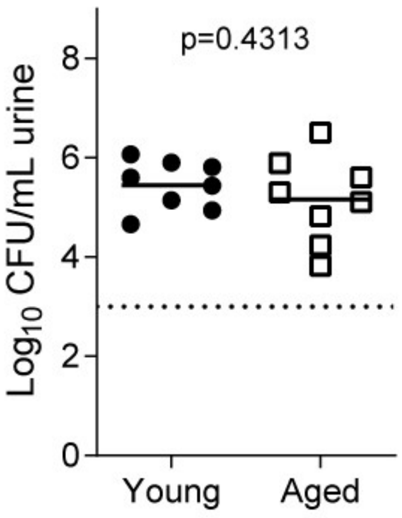


**Fig. S3. Young and aged mice and equally infected by uropathogenic *E. coli* (UPEC).** Urine bacterial titers from young and aged mice 6 hours post infection. Dotted line is limited of detection (y=3).

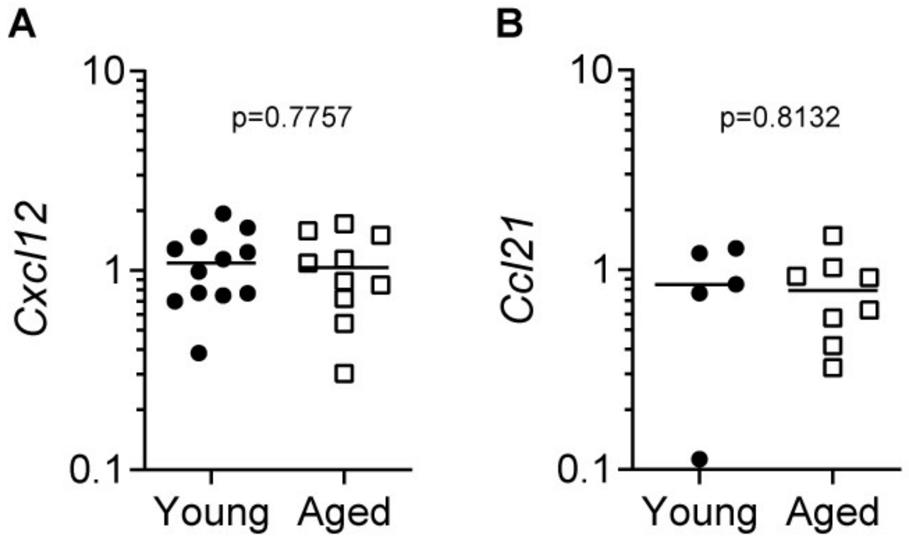


**Fig. S4. *Cxcl12* and *Ccl21* expression are unchanged in bladders from aged mice compared to those from young.** (**A**) Relative *Cxcl12* in bladders from young (n=12) and aged (n=10) mice. (**B**) Relative *Ccl21* expression in bladders from young (n=5) and aged (n=8) mice.
